# PTEX helps efficiently traffic haemoglobinases to the food vacuole in *Plasmodium falciparum*

**DOI:** 10.1101/2022.11.15.516562

**Authors:** Thorey K. Jonsdottir, Brendan Elsworth, Simon Cobbold, Mikha Gabriela, Sarah C. Charnaud, Madeline G. Dans, Molly Parkyn Schneider, Malcolm McConville, Hayley E. Bullen, Brendan S. Crabb, Paul R. Gilson

## Abstract

A key element of *Plasmodium* biology and pathogenesis is the trafficking of ~10% of the parasite proteome into the host red blood cell (RBC) it infects. To cross the parasite-encasing parasitophorous vacuole membrane, exported proteins utilise a channel-containing protein complex termed the *Plasmodium* translocon of exported proteins (PTEX). PTEX is obligatory for parasite survival, both *in vitro* and *in vivo*, suggesting that at least some exported proteins have essential metabolic functions. However, to date only one essential PTEX-dependent process, the new permeability pathway, has been described. To identify other essential PTEX-dependant proteins/processes, we conditionally knocked down the expression of one of its core components, PTEX150, and examined which metabolic pathways were affected. Surprisingly, the food vacuole mediated process of haemoglobin (Hb) digestion was substantially perturbed by PTEX150 knockdown. Using a range of transgenic parasite lines and approaches, we show that two major Hb proteases; falcipain 2a and plasmepsin II, interact with PTEX core components, implicating the translocon’s involvement in the trafficking of Hb proteases. We propose a model where these proteases are translocated into the PV via PTEX in order to reach the cytostome, located at the parasite periphery, prior to food vacuole entry. This work offers a another mechanistic explanation for why PTEX function is essential for growth of the parasite within its host RBC.

**Author summary:** *Plasmodium falciparum* is the causative agent of the most severe form of malaria in humans, where the symptoms of the disease are derived from the continuous asexual replication of the parasite within the human red blood cells (RBCs) it infects. To survive within this niche, the parasite exports hundreds of parasite effector proteins across the vacuole it resides within and into the RBC. About a quarter of the exported proteins appear to be essential during the blood stage but the functions of these proteins largely remain uncharacterised. Protein export is facilitated by an essential protein complex termed the *Plasmodium* translocon of exported proteins (PTEX). Conditional depletion of PTEX’s core components results in rapid parasite death presumably because essential proteins do not reach their functional destination in the RBC and their associated metabolic functions cannot be performed. To uncover what these essential metabolic functions are we knocked down PTEX150, a core component of PTEX. Metabolic analysis of the knockdown parasites indicated that haemoglobin (Hb) digestion was inhibited resulting in a reduction of Hb derived peptides, which serve as an amino acid source for the parasite. We determined that knocking down HSP101, another PTEX core component, also disrupted the Hb digestion pathway. Furthermore, we provide evidence that reduction of Hb digestion might be due to the failure to efficiently deliver early acting Hb digesting proteases to the cytostome, a specialised location where vesicles of Hb are taken into the parasite. PTEX may therefore play a role in delivering Hb proteases to the cytostome.

## Introduction

Malaria is a disease caused by *Plasmodium* parasites and transmitted to humans with the bite of an infected female *Anopheles* mosquito. Malaria remains a major health and economic burden and it is estimated that 627,000 people died of malaria in 2020 (1). The clinical symptoms of malaria are derived from the asexual replicative stage of the parasite, which occurs within the red blood cells (RBCs) the parasite infects (2). During this stage the parasite invades the RBC whereupon it resides within a parasitophorous vacuole (PV); a membranous sac serving to occlude the parasite away from the RBC cytosol. The intracellular parasite is thus encased in two membranes, the parasite plasma membrane (PPM) and the parasitophorous vacuole membrane (PVM) (3). For survival within the infected RBC (iRBC), the parasite employs its own ATP-powered protein conduit at the PVM termed the *Plasmodium* translocon of exported proteins (PTEX) to export parasite effector proteins across the PVM and into the iRBC to establish essential host cell modifications (4–7).

PTEX is 1.6 MDa complex comprising three core proteins: Exported protein 2 (EXP2), PTEX150 and heat shock protein 101 (HSP101). Within this complex, EXP2 exists as a heptamer, which forms the PVM pore and binds directly to PTEX150, another hepatmer that serves as a stable channel for protein export. HSP101 exists as a hexamer and unfolds proteins, which it then extrudes through PTEX150 for transit through EXP2 and into the iRBC (4, 8–10). These three components cannot be knocked out in the human malaria parasite *P. falciparum* or the rodent malaria parasite *P. berghei* and conditional knockdown results in rapid parasite death and a block in protein export across the PVM, indicating PTEX mediated export is essential for parasite survival (4–7, 11–13).

Bioinformatic analyses have helped predict which proteins are likely exported based on the presence of a pentameric motif at their N-terminus called the *Plasmodium* export element (PEXEL) motif and other export related features (14–18). Some exported proteins, however, lack the PEXEL motif, and are referred to as PEXEL negative exported proteins (PNEPs). Since PNEPs lack a signature export motif it is harder to predict their export (19). It is hypothesised that ~25% of the predicted exported proteome (exportome) is essential for *in vitro* blood stage growth but the essential functions remain unclear (20, 21). To date, only one essential function has been assigned to exported proteins, the establishment of the new permeability pathways (NPPs) at the iRBC surface, which help import essential nutrients from the surrounding blood plasma (22–27). However, most exported protein functions identified are not essential for *in vitro* growth and contribute to iRBC rigidity, virulence and immune evasion (28–30).

Here we conditionally knocked down one of PTEX’s core components, PTEX150, and used metabolomics to identify biological processes affected when PTEX’s function is perturbed and thereby the potential function(s) of the essential exportome. We initially anticipated a relative reduction in the levels of molecules known to enter the iRBC via the NPPs but instead discovered an unexpected link between haemoglobin (Hb) digestion and PTEX function. To strengthen this finding, we also observed an association of two major Hb proteases, falcipain 2a (FP2a) and plasmepsin II (PM II) with PTEX. Overall, the data provided in this study suggests that PTEX core components help with efficient trafficking of Hb proteases within the PV space *en route* to the food vacuole where Hb digestion occurs.

## Results

### 2.1 Conditional knockdown of PTEX150 reduces the level of digested Hb

To perturb the function of one of PTEX’s principal components, we used a previously established parasite line where PTEX150 was appended with a triple haemagglutinin (HA) protein tag and a *glmS* riboswitch to conditionally knockdown PTEX150 in the presence of glucosamine (GlcN) (6, 31). Metabolomic analyses were performed on 18, 24 and 30-h post invasion (hpi) parasites synchronised to a 4 h window one cell cycle after the addition of 0.15 mM or 1 mM GlcN to induce knockdown of PTEX150 expression (Fig 1A).

**Fig 1.**
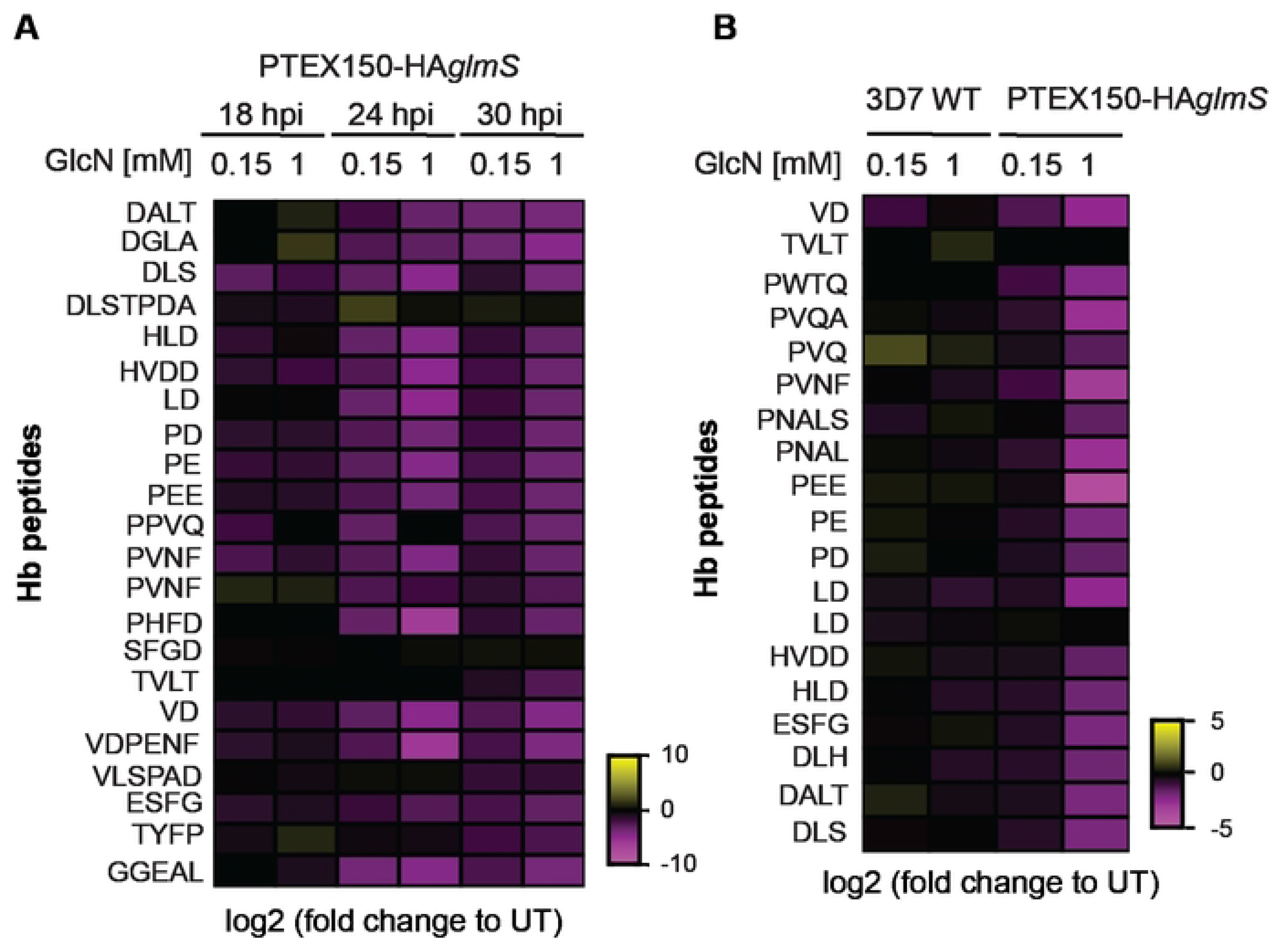
Conditional knockdown of PTEX150 perturbs the Hb digestion pathway. (A) Highly synchronous PTEX150-HA*glmS* trophozoite stage parasites were treated with 0, 0.15 or 1 mM GlcN for one cell cycle to knockdown the expression of PTEX150. The parasites were harvested at 18, 24 and 30-hpi and their metabolites were extracted and fractionated for identification by mass spectrometry. The fold change for 0.15 and 1 mM GlcN compared to untreated (UT) parasites is indicated. The amino acid sequences of the Hb peptides are shown on the left of the heat map. Heat map shows 1 biological replicate completed in technical triplicate. (B) Metabolites from 3D7 WT and PTEX150-HA*glmS* parasite lines were prepared as described for panel A and parasites harvested at 24-hpi. The heat map represents 1 biological replicate of 3 technical replicates, where fold change is shown for 0.15 or 1 mM GlcN compared to untreated. Fold change for both heat maps was calculated using the following formula =log2(GlcN treated/untreated).

Analysis of resultant data revealed that glycolysis and nucleotide metabolism was perturbed in PTEX150 knockdown parasites, indicating cellular homeostasis was dysregulated (S1 Table). This was predominantly observed in parasites 30-hpi, a treatment time at which the PTEX150 knockdown parasites are known to stall (6). These effects are therefore most likely a result of parasite growth arrest rather than a direct involvement of PTEX in these pathways (S1 Table).

The data also revealed that PTEX150 knockdown resulted in a noticeably lower abundance of Hb peptides when compared to untreated samples, indicating that Hb digestion was reduced in these parasites (Fig 1A, S1A Fig, S1 Table). The largest difference in Hb peptides abundance was observed at 24-hpi and 30-hpi in parasites treated with 1 mM GlcN (Fig 1A). However, unlike for glycolysis and nucleotide metabolism, this effect was also observed in parasites treated with 0.15 mM GlcN, which does not significantly affect parasite growth (Fig 1A, S1A Fig) (6). Therefore, reduction of digested Hb is likely due to PTEX150 knockdown and not due to a defect in overall parasite growth. In further support of this, a moderate decrease in certain Hb peptides was also observed at 18-hpi and a noticeable decrease in 24-hpi parasites treated with either 0.15 or 1 mM GlcN, prior to when growth stalls due to PTEX150 knockdown (Fig 1A). Overall, these data indicate that knockdown of PTEX150 specifically reduces Hb digestion.

To corroborate these findings, the experiment was repeated using a single time point and a 3D7 wild-type (WT) parasite control to monitor any effects caused by the GlcN treatment itself on Hb peptide abundance. The parasites were synchronised and GlcN-treated as before using a single time point (24-hpi). This time point was chosen because the majority of Hb digestion occurs 18 – 32-hpi (32, 33). As previously observed, the most dramatic metabolic differences between the lines following PTEX150 knockdown are in the levels of Hb peptides (Fig 1B, S1B Fig, S1 Table). Specifically, the 3D7 WT parasites showed a slight decrease in some Hb peptides when treated with GlcN, but the reduction observed for PTEX150-HA*glmS* expressing parasites was much greater (Fig 1, S1 Fig). Collectively these data indicate that PTEX likely plays a role in the Hb digestion pathway.

### 2.2 Establishing a falcipain 2a knockdown line as a positive control for disturbance to Hb digestion

It was not immediately obvious how knockdown of PTEX reduces Hb peptides and so we postulated that PTEX could either be involved in the uptake of Hb or involved in the trafficking of the early acting falcipain and plasmepsin Hb proteases to the cytostome *en route* to the food vacuole / digestive vacuole (34–36). The cytostome is an invagination of both the PPM and the PVM from which Hb containing vesicles bud off for transport to the food vacuole where the majority of Hb digestion occurs (37, 38). Uptake of Hb and digestion are both important for parasite survival, providing both amino acids (aa) and space for the growing parasite (33, 39–41). Falcipain 2a (FP2a) is one of the early acting Hb proteases (42–44) and due to this, it was chosen for this study to determine if knocking down FP2a expression would mimic the effects observed for PTEX150 knockdown. We appended the C-terminus of the *fp2a* gene (PF3D7_1115700) with a single HA-tag and the *glmS* riboswitch using the CRISPR/Cas9 approach (S2A Fig). C-terminal tagging has not previously been found to affect its trafficking or function (34–36). Furthermore, FP2a is not individually essential for *in vitro* growth (45, 46), therefore knocking it down would not be expected to severely reduce parasite growth.

Correct integration of the tag to the *fp2a* locus was confirmed via PCR (S2B Fig) and western blotting was used to confirm the presence of FP2a-HA*glmS*, where both pro-(54 kDa) and mature (27 kDa) forms of the protease were observed as has been reported previously (43, 47) (S2C Fig). To investigate the level of knockdown, FP2a-HA*glmS* trophozoite stage parasites were treated with increasing concentrations of GlcN for one cell cycle and >90% protein knockdown was observed by western blotting (S2C Fig). Immunofluorescence assays (IFAs) were then used to determine the localisation of FP2a-HA*glmS* within iRBC where it showed diffuse localisation in the parasite cytoplasm (S2D Fig), often with concentrated puncta around the parasite surface, possibly representing the cytostome as has been previously observed (34, 36, 48) (S2D Fig, white arrows). While green fluorescent protein (GFP) tagged FP2a is shown to concentrate in the food vacuole, FP2a-HA*glmS* was found throughout the parasite cytoplasm as previously observed with a native FP2a antibody (34). This is not entirely unexpected, as previous reports of a HA-tagged food vacuole protein, lipocalin (PF3D7_0925900), showed that the HA-tag is degraded upon food vacuole entry (49). The HA-tag of our FP2a-HA*glmS* could therefore also be degraded and not easily detectable inside the food vacuole by IFA.

To assess parasite growth upon FP2a-HA*glmS* knockdown, multi-cycle growth assays were conducted on both 3D7 WT and FP2a-HA*glmS* parasites. Trophozoite stage parasites were treated with increasing concentrations of GlcN over three consecutive cell cycles and harvested at each cycle to measure lactate dehydrogenase (LDH) activity as a proxy for parasite growth (50, 51). As expected, there was no growth reduction observed in the FP2a-HA*glmS* knockdown parasites relative to untreated (0 mM GlcN) parasites or the 3D7 WT control which concurs with previous FP2a knockout studies (52) (S2E Fig).

### 2.3 Knocking down PTEX’s core components, PTEX150 and HSP101, causes a build-up of full-length Hb inside the parasite

To determine if reduced Hb peptides upon PTEX150 knockdown was due to either reduced Hb uptake, or reduced Hb digestion, we completed western blotting on both the PTEX150-HA*glmS* line, and an additional line in which the core PTEX component HSP101 was similarly tagged (HSP101-HA*glmS*) (53). As a control for Hb digestion we utilised the FP2a-HA*glmS* parasite line, as well as 3D7 WT parasites as a negative control.

To complete these assays, tightly synchronised trophozoite stage parasites (~24-hpi) were treated with 0, 0.15 or 1 mM GlcN for one cell cycle and the RBCs were subsequently lysed in 0.09% saponin to remove RBC Hb. The parasite pellets were washed extensively in PBS to remove Hb contamination prior to western blotting (Fig 2A and 2B). As expected, the 3D7 WT control showed minimal changes in Hb levels, whereas following knockdown of PTEX150-HA*glmS*, HSP101-HA*glmS* or FP2a-HA*glmS*, full-length Hb was found to accumulate inside the parasite. This accumulation was found to be inversely proportional to the level of knockdown; a reduction in PTEX150-HA*glmS*, HSP101-HA*glmS* or FP2a-HA*glmS* proteins was associated with an increase in undigested Hb inside the parasite (Fig 2A and 2B). Importantly, this effect was not due to an overall growth defect in the *glmS* lines upon protein knockdown as build-up of full-length Hb was also detected at low GlcN concentrations (0.15 mM) at which parasite growth of *FP2a-HAglmS* (S2E Fig) and PTEX150-HA*glmS* (6) was not substantially perturbed. Although the Hb-build up was only found to be significant for the FP2a-HA*glmS* parasites when comparing % Hb build-up compared to untreated, both PTEX150-HA*glmS* and HSP101-HA*glmS* showed a strong trend for Hb-build up with increased protein knockdown. We were not able to establish a strong knockdown for either PTEX150-HA*glmS* or HSP101-*HAglmS*, which might be why we don’t observe a significant increase in full-length Hb inside. FP2a was almost completely knocked down which is likely why we were able to achieve statistically significant results. We therefore also performed a simple linear regression analysis, where all three parasite lines (PTEX150-HA*glmS*, HSP101-HA*glmS* and FP2a-HA*glmS*) showed significant regression slope for Hb build-up when protein expression was reduced (S3A Fig).

**Fig 2.**
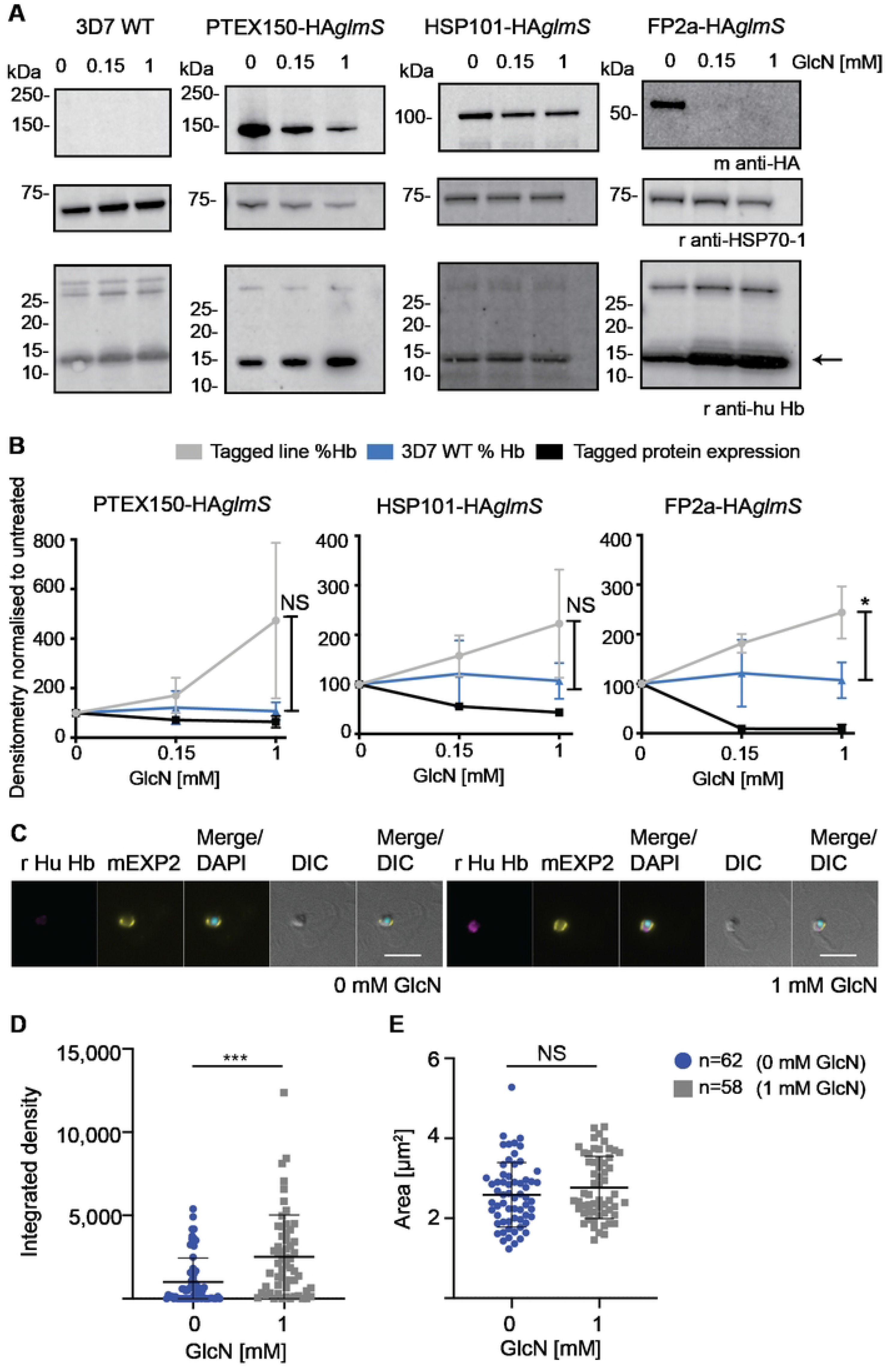
Knockdown of PTEX150 and HSP101 results in build-up of full-length Hb inside the parasite. (A) Trophozoite stage parasites were treated with 0, 0.15 or 1 mM GlcN for one cell cycle, harvested via saponin lysis at ~20-24-hpi and prepared for western blotting. Mouse anti-HA was used to probe for the target protein, rabbit anti-HSP70-1 was a loading control and rabbit anti-human Hb a marker for undigested Hb inside the parasites. Blots are representative of 3 biological replicates. (B) Protein band densitometry of 3 biological replicates was used to indicate the relative levels of Hb in the GlcN-induced knockdown parasites normalised to untreated parasites. 3D7 WT was used as negative control and FP2a-HA*glmS* as a positive control for Hb build-up. The densitometry for Hb was done on the monomer (indicated with an arrow in panel A). The densitometry was adjusted to the loading control and normalised to untreated to visualise changes when treated with GlcN. Error bars= SD. “%” refers to % build-up of Hb compared to untreated and “exp.” refers to protein expression of the knockdown protein following GlcN treatment. Statistical analysis was completed for 1 mM GlcN comparing 3D7 WT to PTEX150, HSP101 and FP2a-HA*glmS* lines using Student’s t test with Welch correction. Only FP2a showed significant increase in Hb levels compared to WT, where * P=0.0255. However, both PTEX150 and HSP101 knockdown showed trend towards Hb build-up inside the parasite when knocked down. (C) Highly synchronous PTEX150-HA*glmS* parasites were treated ± 1 mM GlcN for one cell cycle and 20-24 hpi trophozoites were lysed in saponin and prepared for IFA. Panels show representative figures from 1 biological replicate. DAPI (4’,6-diamidino-2-phenylindole) was used to stain the nucleus. DIC=differential interference contrast. Scale bars= 5μm. (D) For cells from IFA experiments in panel C the integrated density (mean intensity*area) of the parasite, denoted by the boundary of EXP2 signal, was measured in the Hb channel. Statistical analyses were completed using Student’s t test with Welch correction, where the number of cells analysed is indicated on the right-hand side. A significant increase in Hb integrated density was observed when parasites were treated with 1 mM GlcN compared to 0 mM GlcN, where (***) indicates P=0.0001 and error bars=SD. (E) Cells analysed in panel D were of similar size where Student’s t test with Welch correction showed that there was no significant difference between treatments. Middle line represents mean and error bars=SD.

To complement our results, we completed IFAs on saponin-lysed parasites probed with antibodies against full-length Hb (Fig 2C, 2D and 2E). Following 1 mM GlcN treatment of PTEX150-HA*glmS* parasites as described above, full-length Hb was also observed accumulating within the parasite, independent of parasite size, confirming previous results (Fig 2C, 2D and 2E).

As Hb is digested by the parasite, toxic by-products from this process are sequestered as haemozoin crystals, which can be observed by light microscopy. In the absence of Hb digestion, these crystals are therefore not formed (54). We investigated the presence of haemozoin crystals in our PTEX knockdown lines. Specifically, either PTEX150-HA*glmS* or HSP101-HA*glmS* were treated for one cell cycle ± 2.5 mM GlcN to reduce respective protein expression (S3B, S3C and S3D Fig). 2.5 mM GlcN treatment was used because it produces the largest degree of knockdown for *glmS*-tagged proteins without noticeably reducing the growth of non-tagged parasites (6, 53). Both PTEX150-HA*glmS* and HSP101-HA*glmS* knockdown resulted in significantly less haemozoin crystal formation (S3B and S3C Fig) indicating less Hb was being digested which concurs with western blotting and IFA data above (Fig 2). At this level of GlcN, we did observe a significant reduction in parasite size for the PTEX150-HA*glmS* parasites but not for HSP101-HA*glmS* (S3D Fig). Collectively these data demonstrate that upon modest knockdown of the PTEX components PTEX150-HA*glmS* and HSP101-HA*glmS*, parasites still take up Hb. However, the parasites are unable to properly digest Hb indicating that PTEX may be involved in the trafficking of proteases involved in the degradation of Hb, and likely not the process by which the Hb is taken into the parasite.

### 2.4 Generation of FP2a trappable reporter cargoes to investigate FP2a relationship with PTEX

Next, we sought to better understand the relationship between PTEX and Hb digestion. To this end, we generated a series of FP2a reporter constructs to investigate the reporters’ trafficking and interaction with PTEX *en route* to the food vacuole as the protease is thought to traffic to the food vacuole via the cytostome at the PPM/PVM interface (34, 35, 43). Three FP2a reporters of differing lengths were generated, all appended to a nanoluciferase (Nluc) ultra-bright bioluminescence reporter (55), murine dihydrofolate reductase domain (DH) and a triple FLAG (FL) epitope tag. The initial two constructs included the first 120 or 190 aa of FP2a, here referred to as “120 aa” and “190 aa”, respectively (Fig 3A). Previous truncation studies demonstrated that the first 105 aa of FP2a are sufficient for its trafficking to the food vacuole (35), however a construct containing the first 120 aa was more efficiently trafficked to the food vacuole which is why this length was chosen here (34, 35). The longer 190 aa version was made to investigate if an N-terminus longer than 120 aa would provide an even more efficient PPM extraction as this has been shown to be important for exported transmembrane (TM) domain proteins (56). A third reporter was generated as a negative control and contained the N-terminal trafficking region of FP2a but lacked the N-terminal TM domain (Fig 3A) required for entry into the secretory pathway (34, 35) and subsequent trafficking to the food vacuole. This reporter is referred to as “NT” throughout the text (Fig 3A).

**Fig 3.**
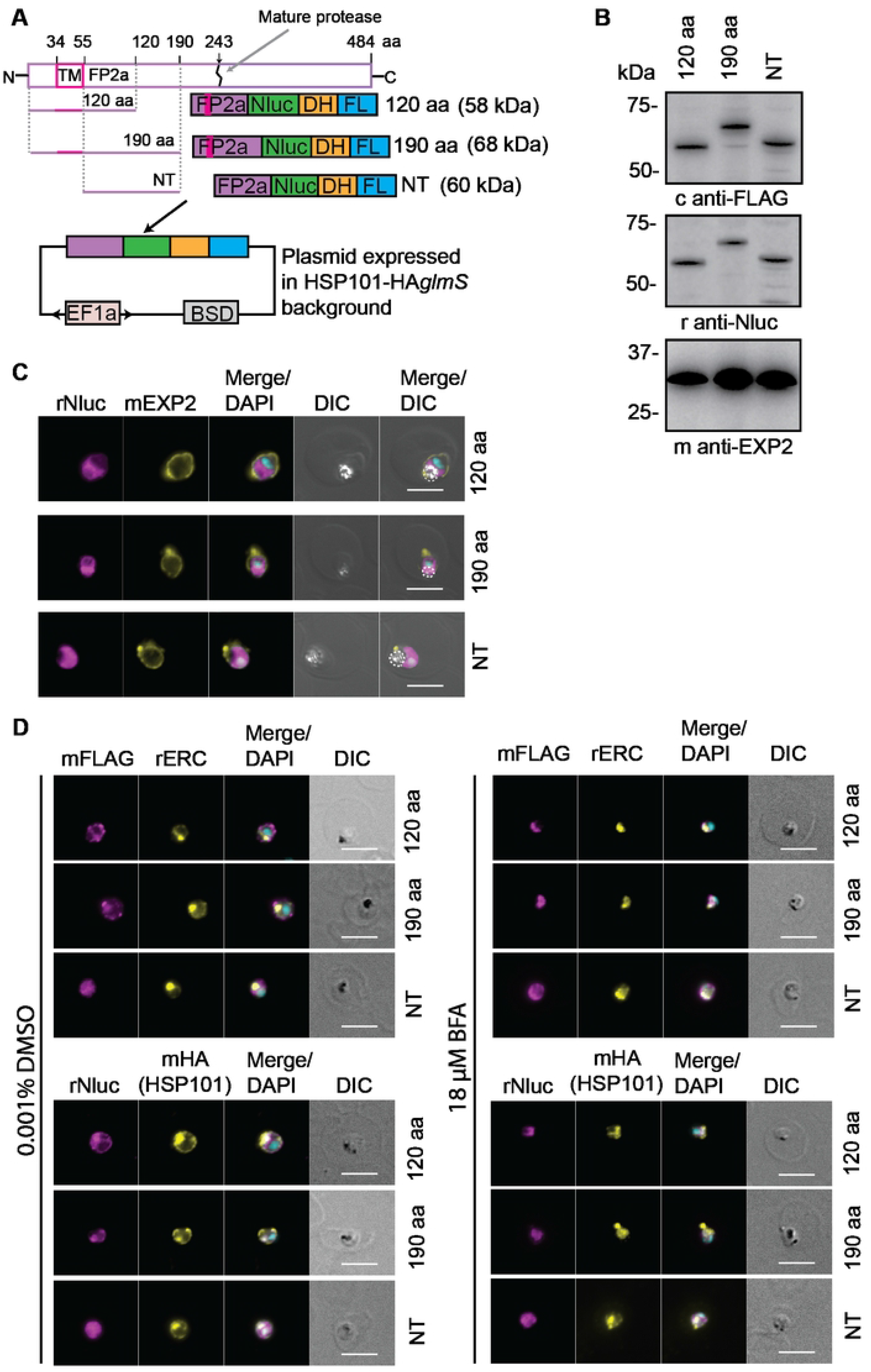
FP2a trappable reporters were generated to study the relationship of FP2a with PTEX. (A) Three FP2a reporters were generated. 120 aa and 190 aa containing the first 120 aa and 190 aa of FP2a respectively. The NT reporter refers to the 190 aa reporter without the N-terminal TM domain and served as a negative control. All FP2a sequences were appended with nanoluciferase (Nluc), murine dihydrofolate reductase (DH) and a 3X FLAG (FL) epitope tag. The complete protein map for FP2a is indicated above for reference, where the reporter only contains the N-terminus sufficient for food vacuole delivery and not the C-terminus necessary for protease activity. The reporters were episomally expressed from a plasmid under the bidirectional EF1a promoter in the HSP101-HA*glmS* parental line and maintained under Blasticidin-S selection via the Blasticidin-S Deaminase (BSD) cassette. (B) Western blot analysis demonstrated that the 120 aa, 190 aa and NT FP2a reporters are expressed and migrate at the correct size indicated in panel A. Rabbit anti-Nluc was used to detect the Nluc tag and chicken anti-FLAG to detect the FL tag. Mouse anti-EXP2 was used as a loading control. (C) Immunofluorescence assays show that the 120 aa and 190 aa FP2a reporters are trafficked to the food vacuole. The NT FP2a reporter displayed diffuse labelling within the parasite as expected. Rabbit anti-Nluc was used to detect the reporter, mouse anti-EXP2 was used as a PVM marker and haemozoin crystals in the DIC show food vacuole localisation (indicated with a white dotted line). (D) Late ring/early trophozoites were treated ± 18 μM BFA to inhibit ER to Golgi protein secretion. The 120 and 190 aa FP2a reporters were both trapped in the ER upon BFA treatment as expected and no changes were observed for the NT reporter. Rabbit anti-ERC was used as an ER marker and mouse anti-HA to label for HSP101, which also traps in the ER upon BFA treatment (53). DAPI was used to stain the nucleus. Scale bars = 5 μm.

By adding an antifolate ligand such as WR99210 (WR) we can stabilise the globular domain of DH and thereby inhibit the reporter proteins from unfolding. The DH domain has been previously utilised to study both mitochondrial protein import (57) and protein export in malaria parasites (56, 58, 59). PTEX requires cargo to be unfolded prior to export (58) and therefore if FP2a is trafficked via PTEX, the 120 and 190 aa reporters should become trapped in PTEX when inhibited from being unfolded via addition of WR as previously observed for exported protein reporters (56, 59).

All reporters were episomally expressed in the HSP101-HA*glmS* background, under the *ef1a* promoter (Fig 3A). They were successfully detected by western blotting, where they migrated at the expected sizes (Fig 3B). Additionally, IFA analysis confirmed that the 120 and 190 aa reporters were trafficked to the food vacuole where they co-localised with the haemozoin crystals whilst the NT reporter displayed a diffuse signal within the parasite cytoplasm, as expected (Fig 3C). The 190 aa reporter appeared to be more concentrated at the food vacuole than the 120 aa reporter indicating that the longer N-terminus could enable more efficient trafficking to the food vacuole (Fig 3C). Brefeldin A (BFA), which blocks protein secretion from the ER to the Golgi (60, 61), was also used to confirm that the reporters were secreted from the ER as happens for the native protease (34, 35). In the presence of BFA the 120 and 190 aa reporters were retained within the ER whilst no changes were observed for the distribution of the NT reporter as expected (Fig 3D). Overall, these data confirm that the reporters are expressed and appropriate for use in subsequent PTEX trapping experiments.

### 2.5 The FP2a 120 aa reporter displays increased co-localisation with EXP2 when inhibited from unfolding

To determine if FP2a associates with PTEX, the 120 aa, 190 aa and NT reporters were trapped using WR and IFAs were performed to detect co-localisation with PTEX components at the PVM. Tightly synchronised (4 h window) ring stage parasite cultures were treated ± 10 nM WR and harvested as trophozoites (24-28-hpi) for IFA. Rabbit anti-Nluc was used to visualise the FP2a reporters and mouse anti-EXP2 served as both a PTEX and PVM marker (Fig 4A). The 120 and 190 aa reporters displayed some labelling around the parasite surface/PVM, which was expected as early acting Hb proteases are known to traffic there for loading into the cytostome prior to delivery to the food vacuole (34–36). To quantify the co-localisation of reporters with EXP2 ± WR, Pearson’s coefficients of the proteins were measured (Fig 4B). The 120 aa co-localised significantly more with EXP2 when treated with WR indicating that more cargo was trapped at the parasite periphery when rendered unfoldable and therefore potentially trapped in PTEX (Fig 4A and 4B). However, the 190 aa and NT reporters did not show a significant difference ± WR treatment (Fig 4A and 4B). Since the 120 and 190 aa reporters are identical except for their length, it is likely that length is influencing trapping efficiency and that shorter cargo is more easily trapped (discussed in later sections).

**Fig 4.**
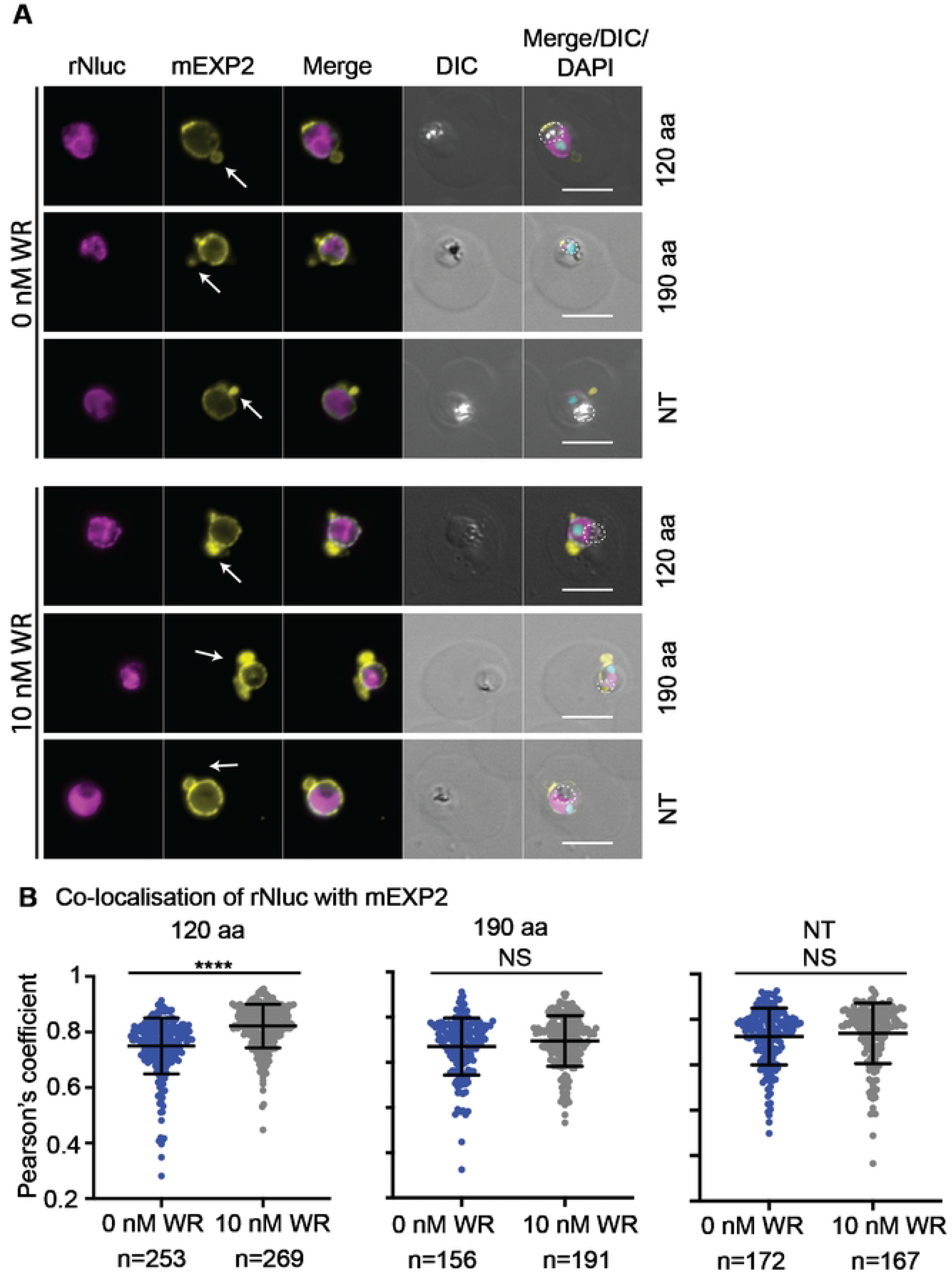
The FP2a 120 aa reporter shows significantly more association with EXP2 upon WR trapping. (A) Parasites expressing the 120 aa, 190 aa and NT FP2a reporters were synchronised and treated ± 10 nM WR at ring stage to prevent unfolding of the reporters to see if they would become trapped at the parasite periphery with PTEX. When treated with WR, more 120 aa reporter was observed at the PPM/PVM indicating its trafficking was affected. No substantial difference was observed for 190 aa or NT reporters. Shown are representative images for 3 (120 aa) and 2 (190 aa and NT) biological replicates. Rabbit anti-EXP2 was used as a PVM marker, rabbit anti-Nluc to detect the reporters and DAPI to stain the nucleus. Scale bars= 5 μm. Arrows point to PVM loops. (B) Pearson’s coefficient was used to measure the co-localisation of the FP2a reporters (Nluc) and EXP2. There was significantly more co-localisation of 120 aa with EXP2 when treated with WR as determined by student’s t test with Welch correction but no significant difference was observed for the 190 aa or NT reporters. (****) Indicates P <0.0001, n= cells analysed (pooled from 3 (120 aa) or 2 (190 aa and NT) biological replicates), middle line on the graph represents mean and error bars=SD. Each dot on the graph represents a cell analysed.

Interestingly, even though the 120 aa cargo co-localised with EXP2, it was never observed inside EXP2 loops (Fig 4A, white arrows) which we commonly observe for the exported reporter Hyp1-Nluc-DH upon WR trapping (59, 62). These loops are thought to represent accumulated trapped cargo unable to be transported by the PTEX complex at the PVM (59). The absence of the FP2a reporter cargo from these loops could indicate that it might not be trapped at the PVM, but instead at the PPM. We sought to further investigate the putative interaction between PTEX and FP2a using co-immunoprecipitation assays (Co-IPs).

### 2.6 Both the 120 aa and the 190 aa reporters associate with PTEX core components, but this association is diminished when reporters are inhibited from unfolding

To investigate the interaction between HSP101 and FP2a, Co-IPs were performed with the three FP2a reporters (NT, 120 aa, 190 aa). We also employed an exported protein reporter, Hyp1-Nluc-DH-FL or “Hyp1”, as a positive control for PTEX interaction (53). Synchronised ring stage parasites were treated ± 10 nM WR and trophozoite stage iRBCs (~28-32-hpi) were isolated from uninfected RBCs (uRBCs) by magnet purification. This time point was chosen because it was when the FP2a and Hyp1 reporter proteins were optimally expressed with less degradation (S4 Fig) (53). Parasite pellets were lysed in modified RIPA buffer and the resultant lysate was incubated with anti-HA agarose beads to immunoprecipitate (IP) HSP101-HA*glmS* and its interacting proteins for visualisation by western blotting (Fig 5A).

**Fig 5.**
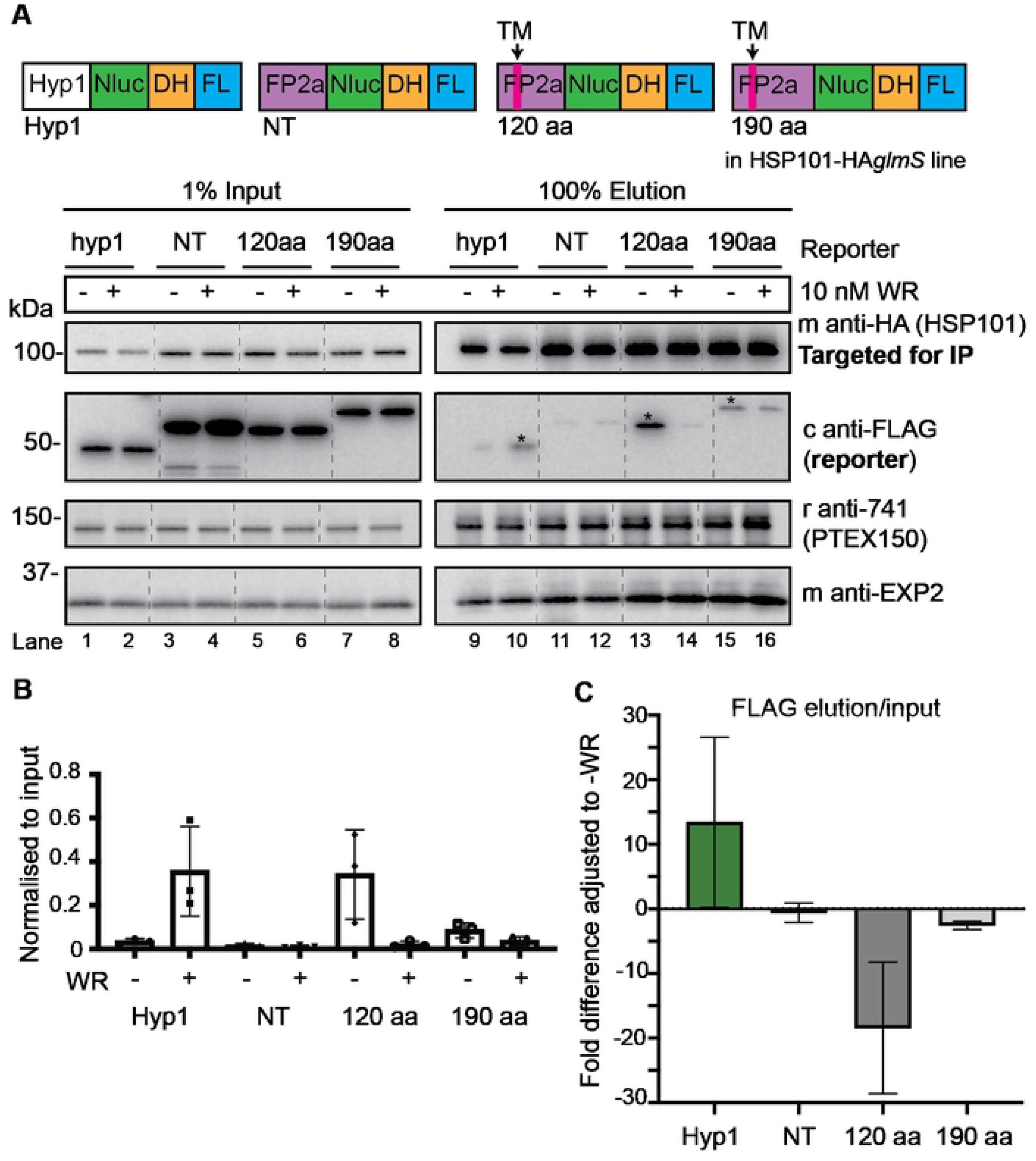
Both the 120 aa and the 190 aa FP2a reporters interact with HSP101 and this interaction is weaker when reporters are inhibited from unfolding via addition of WR. (A) Ring stage HSP101-HA*glmS* parasites episomally expressing Hyp1, NT, 120 aa, or 190 aa reporters were treated ± 10 nM WR and harvested at trophozoite stage (~28-32 hpi). Anti-HA IgG beads were used to IP HSP101 and its interacting proteins. Input shows 1% protein lysate used for the assay, and elution refers to protein bound to the HA beads/HSP101-HA*glmS* indicating their interaction with HSP101. Hyp1 was used as positive control and NT as a negative control. Mouse anti-HA was used to probe for HSP101, chicken anti-FLAG for the reporter, rabbit anti-741 for PTEX150 and mouse anti-EXP2 for EXP2. Blot is representative of 3 biological replicates. The asterisks indicate stronger signal when comparing the ± WR treatment for respective reporter as determined by densitometry of all biological replicates. (B) The interaction of reporters with HSP101 was graphed, where FLAG elution was adjusted to input. Each dot represents 1 biological replicate. (C) The interaction of reporters with HSP101, where the fold difference of the FLAG elution/input was adjusted to untreated (-WR). Error bars= SD from 3 biological replicates.

As expected, the Hyp1 reporter showed an increased association with HSP101 when treated with WR reflective of the folded cargo becoming trapped in PTEX under these conditions (53, 59) (Fig 5A, lanes 9 and 10, 5B and 5C). The NT reporter showed a minimal level of association with HSP101 (Fig 5A, lanes 11 and 12, 5B and 5C) as expected. Importantly, both the 120 and 190 aa reporters showed an enhanced association with HSP101 in the absence of WR, demonstrating that FP2a does indeed interact with HSP101 (Fig 5A, lanes 13 – 16, 5B and 5C). It should be noted that a marked difference was observed between the retention of the 120 and 190 aa reporters (Fig 5A lanes 13 and 14 vs. lanes 15 and 16) with HSP101, whereby substantially more of the 120 aa reporter was bound by HSP101. This concurs with the greater trapping efficiency of the 120 aa reporter at the parasite periphery with EXP2 as visualised by IFA (Fig 4).

It was anticipated that in the presence of WR, the reporters would be more strongly associated with HSP101 if they were trapped in PVM-resident PTEX, but instead the association of the reporters with HSP101 was reduced (Fig 5A, lanes 13 – 16, 5B and 5C). These data therefore imply that WR treated 120 and 190 aa reporter proteins might in fact be trapped in the PPM prior to PTEX association in the PV as has been observed for PNEPs (56). This explains why we did not observe the FP2a reporters in PVM loops by IFA (59) as mentioned above.

Association of the FP2a reporters was also confirmed for PTEX150 and EXP2 by Co-IP using either rabbit anti-PTEX150 (r942) (S5A Fig) or rabbit anti-EXP2 (r1167) (S5B Fig) antibodies. PTEX150 showed a similar association with FP2a cargo as HSP101 (S5A Fig, lanes 13 – 16) and a weak association was observed for the cargoes with EXP2 (S5B Fig, lanes 13 – 16). Densitometry measurements showed a similar trend for the association of Hyp1 and FP2a reporters with both PTEX150 and EXP2 when compared with HSP101 IPs (S6 Fig). Overall, these assays helped to confirm that FP2a interacts with all PTEX core components, as was suggested by our metabolomics and Hb digestion/build-up assays.

### 2.7 The FP2a 120 aa reporter becomes trapped at the PPM with the N-terminus facing into the PV compartment when inhibited from unfolding

Given that the 120 aa reporter cargo displayed an increased co-localisation by IFA with EXP2 when inhibited from unfolding (Fig 4B) but showed reduced association with PTEX components when trapped with WR (Fig 5 and S5 Fig), we utilised proteinase K protection assays to determine which iRBC compartment the reporter was being trapped in. For the assay, synchronised ring stage parasites were treated ± 10 nM WR, prior to harvesting at the trophozoite stage (~24-hpi) via magnet purification. Isolated trophozoite-containing RBCs were treated with equinatoxin-II (EQT) to form pores in the iRBC membrane (63) and release soluble exported proteins (Fig 6A, lanes 1 and 6, SN). The subsequent iRBC pellet was then divided into four fractions that were untreated, treated with proteinase K, or subjected to differential lysis conditions and proteinase K treatment. Saponin was used to lyse the parasite’s PVM, releasing the PV contents (Fig 6A, lanes 4 and 9) and Triton X-100 (TX-100) was used to lyse all membranes (Fig 6A, lanes 5 and 10). Specific antibodies were then used as markers for each lysed compartment, where GBP130 was used as a soluble exported protein marker (59, 64), SERA5 as a soluble PV marker (65) and GAPDH as a marker of the parasite cytoplasm.

**Fig 6.**
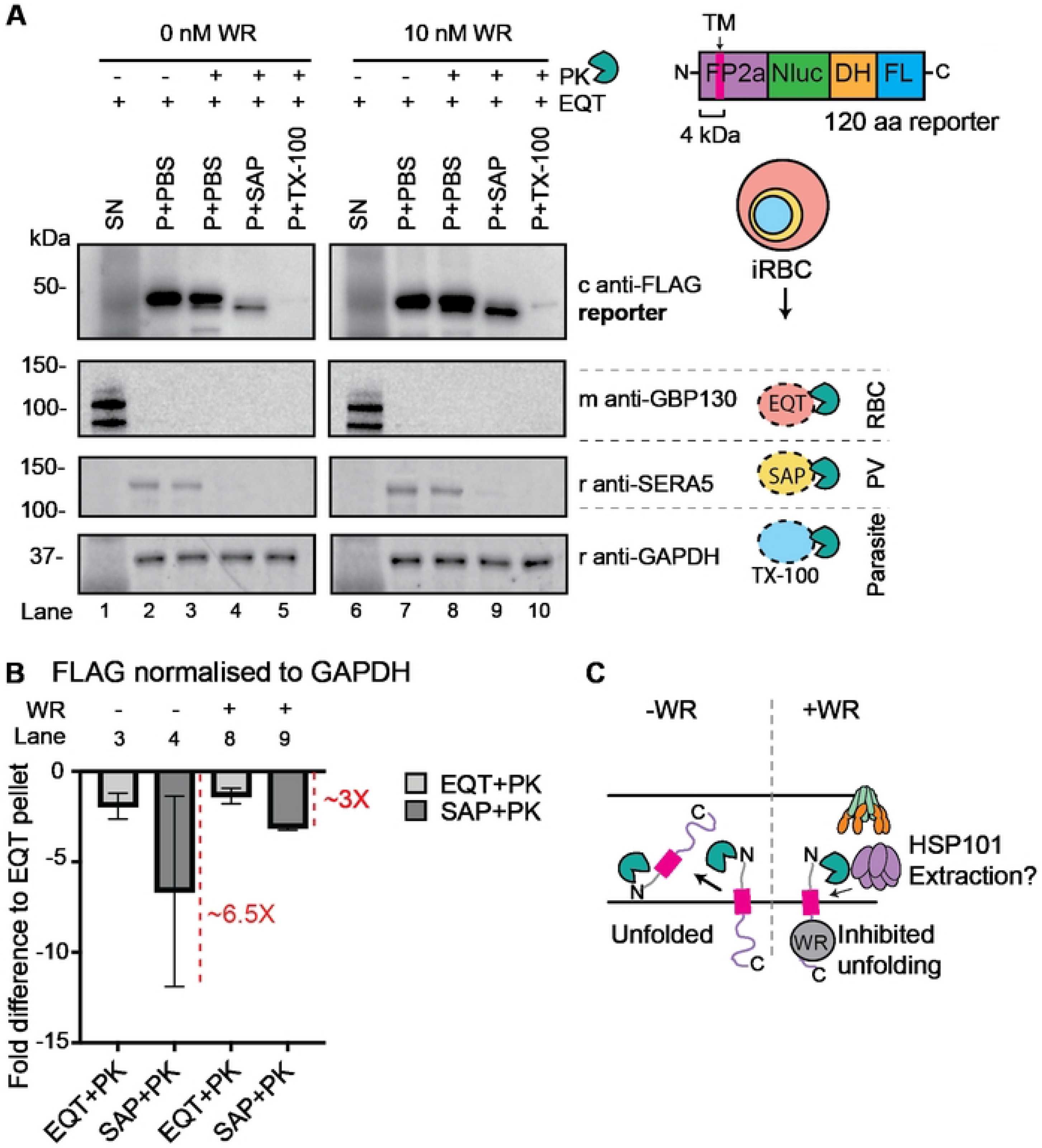
Proteinase K protection assay shows that FP2a needs to be unfolded before crossing the PPM with its N-terminus first. (A) Ring stage parasites were treated ± 10 nM WR to inhibit the 120 aa FP2a reporter from unfolding. Trophozoite stage parasites were magnet purified and used for proteinase K (PK) protection assays. EQT was used to permeabilise the iRBC membrane, saponin (SAP) to lyse the PVM, and TX-100 to lyse all membranes. A 3.8 kDa shift was observed in the reporter size when the PV was lysed and treated with PK (lanes 4 and 9), which matches the predicted size of the 34 aa (4 kDa) N-terminus preceding TM domain (measured in Odyssey LICOR). FP2a reporter appears to be more protected against PK degradation when treated with WR vs. non-treated, indicating trapping at the PPM not PVM. Chicken anti-FLAG was used to probe for the FP2a reporter, GBP130 is a soluble exported protein marker, SERA5 is a PV marker and GAPDH is a parasite cytoplasm marker. Blot is representative of 3 biological replicates, which showed a similar trend, SN= supernatant, P= pellet. (B) The fold difference for protection of the FP2a reporter (FLAG) was measured by densitometry, where the FLAG signal was adjusted to GAPDH and the fold difference measured by adjusting the EQT + PK and SAP + PK signal to EQT + PBS. The fold difference (X) compared with untreated is indicated with a red dotted line on the graph. Error bars = SD from 3 biological replicates. (C) A suggested model derived from the PK assay results. When parasites were not treated with WR more reporter cargo was degraded indicating that when inhibited from unfolding the reporter was trapped at the PPM. This likely hindered extraction into the PV potentially facilitated by HSP101 as suggested for TM exported proteins.

By analysing the distribution of FP2a bands across all treatments, it was possible to infer the orientation and localisation of the 120 aa reporter protein. Firstly, the reporter protein was not degraded in the presence of PBS and proteinase K, indicating no part of the reporter resided within the iRBC cytoplasm, which concurs with our IFA data (Figs 6A, lanes 3 and 8 and 4A). Concordantly, when the parasites were treated with saponin and proteinase K, the FP2a reporter migrated 3.8 kDa smaller than the full-length reporter (Fig 6A, lanes 4 and 9). This 3.8 kDa fragment is proportional to the segment of the reporter protein that precedes the N-terminal TM domain (4 kDa) suggesting that FP2a is oriented with its N-terminus facing into the PV lumen, and the C-terminus is in the parasite cytoplasm (Fig 6C). The FP2a reporter was completely degraded in the presence of TX-100 indicating that the parasite membranes were completely permeabilised (Fig 6A, lanes 5 and 10). GAPDH was not degraded, likely because the protein is highly abundant and proteolytically resistant.

Following WR treatment, the FP2a reporter parasites that were lysed with saponin and treated with proteinase K revealed that the reporter was more resistant to degradation than it was in the absence of WR (Fig 6A and 6B, lanes 3 and 4 vs. lanes 8 and 9) suggesting that FP2a needs to be unfolded before crossing the PPM and entering the PV compartment. This demonstrates that the WR trapping occurs at the PPM interface prior to PTEX association within the PV, which agrees with our IP data where FP2a shows less interaction with PTEX components when treated with WR (Fig 5 and S5 Fig).

### 2.8 PTEX150 co-precipitates native plasmepsin II Hb protease and PTEX150 knockdown affects protease trafficking

To study the relationship between PTEX and a native Hb protease, we tagged the early acting Hb protease plasmepsin II (PM II, PF3D7_1408000) with mScarlet as described for FP2a-HA*glmS* above (S7A Fig), however, the background line used for transfection of this line was PTEX150-HA*glmS*. Integration was confirmed by PCR (S7B Fig). A single band of 75 kDa was detected for PM II-mScarlet by western blotting, where the tag accounted for approximately 27 kDa (Fig 7A). This agrees with previously published results where only the pro form of the protease was detected and not the mature cleaved protease (66). Live cell microscopy confirmed the localisation of PM II-mScarlet during ring and trophozoite stage, where the protease co-localised with the haemozion crystals (Fig 7B, white dotted line) as expected. To determine if PTEX is involved in PM II trafficking, we knocked down PTEX150-HA*glmS* expression at the trophozoite stage and harvested cells for microscopy at the following late ring stage (~20-hpi). We observed a significant accumulation of PM II-mScarlet at the parasite periphery when PTEX150 was knocked down (crescent/circle) compared to a peripheral ‘dot’ when PTEX150 was normally expressed (Fig 7C and 7D). While this is qualitative evidence for a role for PTEX150 in PM II-mScarlet trafficking to the food vacuole, some observations could merely represent PM II-mScarlet during its normal trafficking to the cytostome via the parasite periphery. To support that our observations are indeed due to an interaction with PTEX150, we targeted PTEX150-HA*glmS* for IP using anti-HA agarose beads as previously described. A strong association was seen between PTEX150-HA*glmS* and PM II-mScarlet (Fig 7E). Positive control antibodies against HSP101 and EXP2 also revealed that they too associated with PTEX150 as expected, while rabbit anti-GAPDH was used as a negative control and showed no association (Fig 7E). 3D7 WT parasites were used to detect any non-specific antibody binding in the assay and showed no interaction with PTEX150 or other PTEX components. Overall, this assay confirmed a specific association between PTEX and a native Hb protease (PM II-mScarlet) and agrees with our FP2a reporter data.

**Fig 7.**
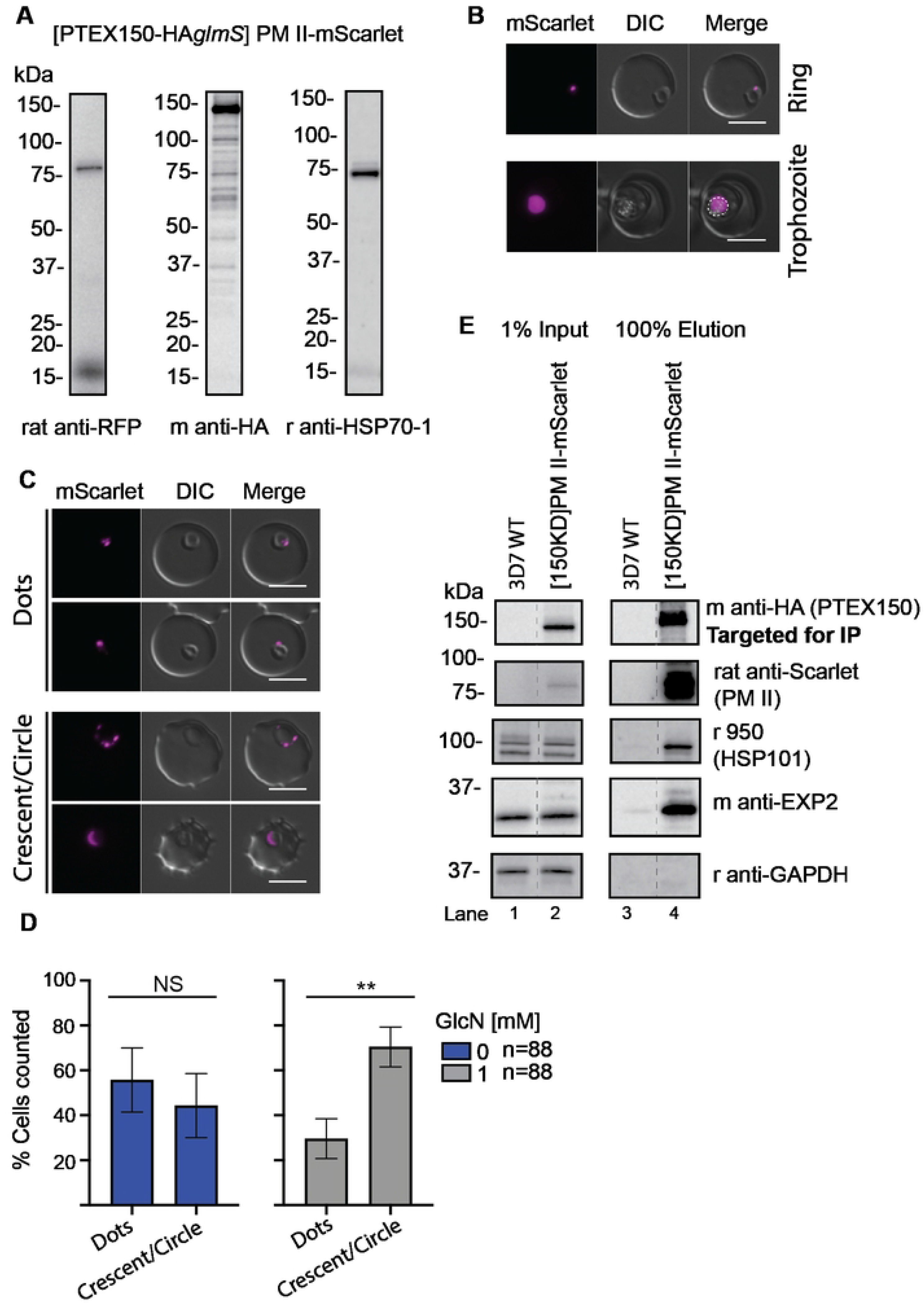
The PM II-mScarlet fusion protein in the PTEX150-HA*glmS* parental line accumulates at the parasite periphery when PTEX150 is knocked down and co-precipitates with PTEX150. (A) Western blot was used to confirm the correct size of PM II-mScarlet, detected by rat anti-RFP (red fluorescent protein), which detects the mScarlet tag. Mouse anti-HA was used to detect the PTEX150-HA*glmS* of the parental line and rabbit anti-HSP70-1 was used as a loading control. Only the pro form of the protease was detected. (B) Live cell microscopy was used to visualise the localisation of the mScarlet (PM II) in both ring and trophozoite stage parasites, where the tag was observed inside the parasite and overlapping with the food vacuole (haemozoin crystals), which are indicated with a dotted white line. Scale bars= 5 μm. (C) PM II-mScarlet parasites were treated ± 1 mM GlcN at trophozoite stage and harvested at late ring/early trophozoite stage (~20 hpi) for live cell microscopy. The mScarlet signal was subsequently categorised as either ‘dot’ or ‘crescent/circle’ around the parasite periphery. Panel shows representative images for each category, the experiment was completed in 3 biological replicates. Scale bars=5μm. (D) Graphs show the percentage of parasites with dots vs. crescent/circular labelling from the experiment in panel C. Statistical analysis was completed using Student’s t test with Welch correction. n = total number of cells counted per condition with 3 biological replicates combined. Error bars= SD for 3 individuals counting. (**) Indicates P= 0.0048. (E) Anti-HA IgG beads were used to IP PTEX150 and its interacting proteins. EXP2 and HSP101 were used as positive control for interaction, GAPDH as a negative control and 3D7 WT as a negative control for non-specific interaction with beads. Blot is representative of 2 biological replicates.

## Discussion

This study sought to elucidate why PTEX is essential by determining which metabolic pathways were perturbed due to the failure to export proteins into the iRBC compartment when the protein components of PTEX were knocked down. Hundreds of parasite proteins are exported into the iRBC, but we do not know what most of these proteins do and why some of them are essential. Metabolomic analyses revealed that knockdown of PTEX150 caused a slight dysregulation of some metabolite levels, for example those involved in glycolysis and nucleotide metabolism, however this was also observed in the 3D7 WT control indicating a non-specific effect from the GlcN treatment. The only clearly defined effect of PTEX150 knockdown was upon Hb digestion, possibly linking the PTEX machinery to this essential catabolic process that provides the parasite with amino acids and helps maintain osmotic stability of the iRBC (33, 39–41). Using several approaches, we were able to determine a physical connection between PTEX and two major early acting Hb proteases, FP2a and PM II. It is possible that these Hb proteases might rely on PTEX components as they transit via the PPM *en route* to the cytostome (34–36).

Modest knockdown of PTEX150-HA*glmS* with 1 mM GlcN reduces PTEX150-HA*glmS* expression to ~60% of normal levels (in our study) and arrests growth at the late ring / early trophozoite stage in the cycle following addition of GlcN (6). As the degree of growth inhibition would also likely produce many metabolic defects, we also used 0.15 mM GlcN, which does not arrest parasite growth (6). This treatment also resulted in a reduction in Hb digestive peptides and western blotting and microscopy analysis demonstrated the reduction was likely due to diminished capacity to digest Hb rather than a defect in total Hb uptake. This contradicts results presented in a recent study by Gupta *et al.*, where they observed less Hb uptake in HSP101 knockdown parasites (67). Their study was however performed differently to ours whereby HSP101 expression was knocked down from late trophozoite stage of the first cycle to the early schizont stage of the next cycle (42-44-hpi). This could result in the HSP101 knockdown parasites stalling at the early trophozoite stage of the second cycle and therefore not being able to take up as much Hb as the untreated control parasites which had continued growing and transitioned into to the schizont stage. This trophozoite stalling effect due to functional inactivation of HSP101 has been previously noted using the same HSP101 knockdown line (5).

To determine if PTEX knockdown was inhibiting the delivery of Hb proteases to cytostomal vesicles we utilised FP2a reporter cargo constructs, which allowed us to study the interactions between PTEX and this early acting Hb protease. The FP2a 120 aa and 190 aa reporters were directly co-precipitated with PTEX core components, although much stronger with HSP101 and PTEX150 than EXP2. If the N-terminus of the FP2a reporter that was poking into the PV (34 aa) was interacting with the PTEX complex, it likely associates more with HSP101 and PTEX150 than EXP2, which is further away (Fig 8). The slightly weaker association with EXP2, that was also observed for the positive control Hyp1, could be due to IP efficiency. EXP2 has a dual function; as a channel in the PTEX complex and as a nutrient channel on the PVM (13, 68). This dual function could therefore influence the co-IP of EXP2 with FP2a and Hyp1, as the pool of nutrient related EXP2 is not associated with PTEX functioning. Importantly, each assay co-precipitated the other members of the PTEX suggesting involvement of the full PTEX complex in protease binding (Fig 5 and S5 Fig) and a similar trend was observed for HSP101, PTEX150 and EXP2 IPs with reporter cargoes (Fig 5B, 5C and S6A and S6B Fig). Importantly, this association was also observed for a native early acting Hb protease, PM II, suggesting PTEX’s interaction with FP2a reporters reflected genuine trafficking interactions (Fig 7E). It should be noted that conditionally inhibiting the function of HSP101 did not reduce the proteolytic processing of native PM II (5) that normally occurs in the food vacuole facilitated via FP2 and FP3 (69). Despite the knockdown of PTEX, some Hb peptides were still produced and some haemozoin crystals did form. This indicates that although PTEX was not essential for the trafficking of PM II (5), the translocon might rather be important for efficient the delivery of Hb proteases to the cytostome. However, given the modest level of knockdown achieved in this study the remaining levels of PTEX expression could be sufficient to sustain protease trafficking and Hb digestion. Although reduction in Hb digestion following PTEX knockdown probably contributes to some growth arrest, the dysregulation of other metabolic pathways might also play a greater role in reducing parasite growth, however, our analysis does not indicate what these other pathways are.

**Fig 8.**
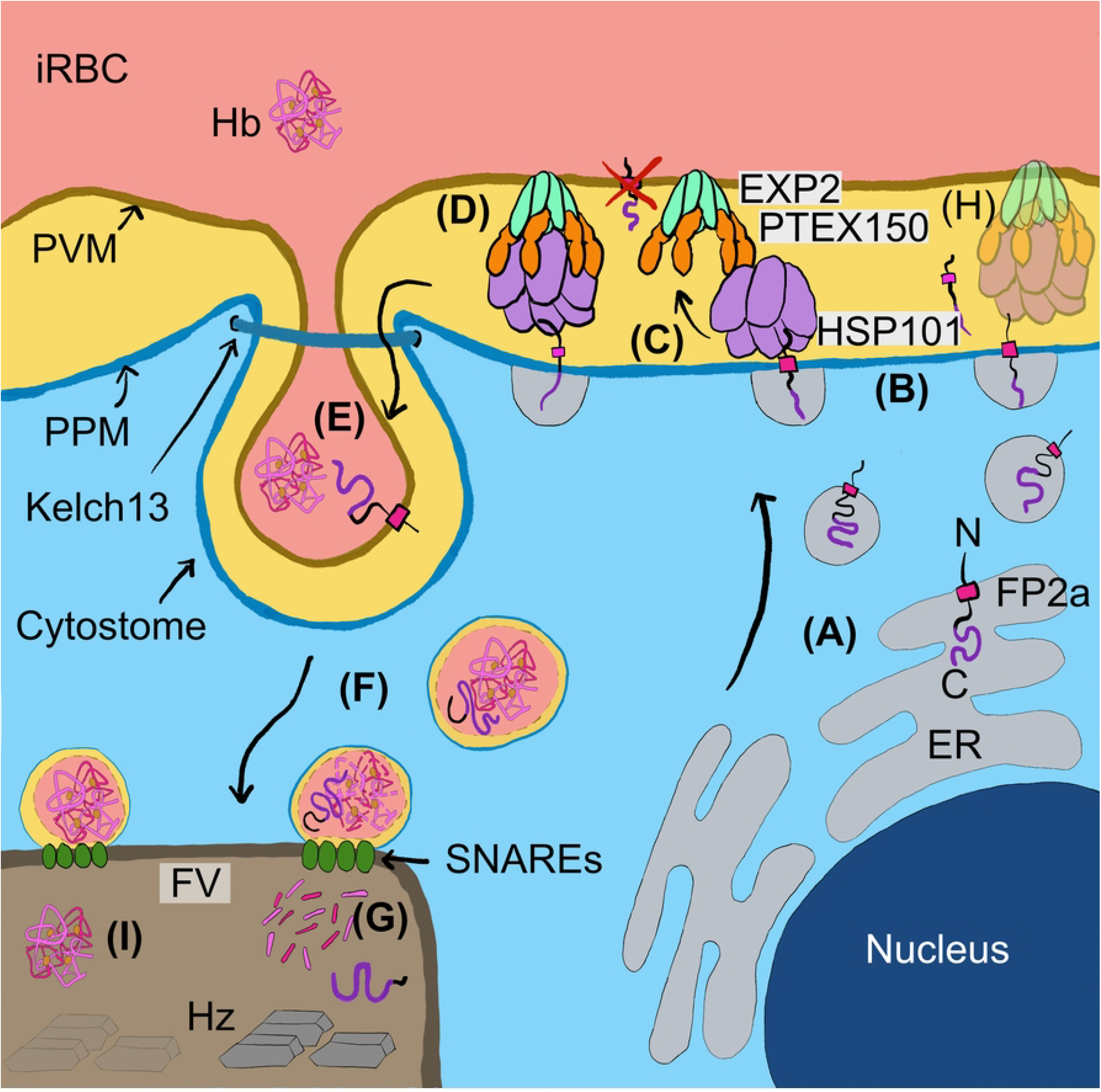
Model depicting newly identified steps in the FP2a trafficking pathway to the food vacuole. By integrating the data from this study, we propose a new model for the trafficking of FP2a, and potentially other Hb proteases, to the food vacuole. (A) The FP2a protein enters the secretory pathway and travels from the ER to the PV, (B) where the protein is deposited in the PPM via its TM domain (pink), in an N- to C-terminal direction. (C) HSP101 could help extract FP2a from the PPM and into the PV as proposed for TM exported proteins and then interact with the rest of PTEX, or (D) all the PTEX core components could participate in FP2a extraction from the PPM. FP2a is then escorted to the cytostome, which likely involves additional sorting in the PV by an unknown mechanism. (E) The proteolytic domain (purple) of FP2a is likely placed with the C-terminus inside the cytostome for easy access to the Hb, which is taken up via the cytostome formed by invagination of the PVM and PPM. (F) FP2a is then transported to the food vacuole (FV) together with Hb. (G) Inside the FV the Hb is digested into smaller peptides, releasing haem, which is sequestered into haemozoin crystals (Hz). (H) Reduced PTEX expression results in accumulation of Hb proteases (as shown with PM II-mScarlet) at the parasite periphery. (I) This would subsequently lead to accumulation of full-length Hb inside the FV and reduced levels of haemozoin crystal formation. Figure is partially based on Hb trafficking models provided by (32, 38). A Red cross is drawn over FP2a incorporation into the PVM by PTEX, as this is likely not possible since PTEX does not have lateral gate to facilitate this.

As FP2a has been shown to traffic to the cytostomal vesicles via the parasite periphery (34, 35) we investigated if PTEX may associate with the FP2a reporters at the PPM. Here we demonstrated that the FP2a reporters likely need to be unfolded prior to crossing the PPM, like exported proteins containing TM domains (10, 56, 70). We showed this by appending an inducibly unfoldable DH domain to the reporters, which in its unfoldable form was more strongly localised to the parasite periphery. Our proteinase K protection assays further indicated the direction of insertion into the PPM likely followed an N- to C-terminal direction. Immunoprecipitation assays additionally showed that PTEX was in contact with the 120 aa and 190 aa FP2a reporters and with the native PM II protein, suggesting PTEX could help extract these proteins into the PV lumen. HSP101 has previously been suggested to facilitate the extraction of TM exported proteins from the PPM (71). An explanation as to how these proteases are trafficked to the cytostome via PTEX is that they are transferred into the PVM via PTEX, which is anchored to the PVM via EXP2, and then incorporated into the innermost membrane of the cytostomal vesicles. Although this is an attractive explanation for how PTEX assists Hb proteases to enter the cytostome, further experiments did not strongly support this scenario and PTEX does not have a lateral gate to facilitate the transfer of proteins to the PVM (9, 72, 73) (Fig 8, red x symbol).

Rather from our data it appears that the Hb proteases are not translocated through PTEX into the PVM since the FP2a reporters containing the DH domain did not co-localise to the PVM loops with EXP2 in the presence of WR as has been shown previously for export-blocked PEXEL-DH reporter proteins (59). Secondly, the 120 aa FP2a reporter was protected from protease degradation when the iRBC was permeabilised with EQT but not when the PVM was lysed with saponin indicating the protease’s short 34 aa N-terminal region upstream of its TM domain was projecting into the PV. The fact that the FP2a reporter was more strongly degraded in the absence of WR indicates that PTEX probably only extracts the reporter as far as the PV and not beyond (Fig 8). If PTEX could capture FP2a reporters and deliver them to the cytostome before translocating them into the cytostome lumen for uptake into vesicles, PTEX would be expected to concentrate at the cytostome, which might appear as one to a few distinct puncta at the parasite periphery (36). We observed these concentrated puncta for FP2a (S2D Fig) but we never observed a particular concentration of EXP2 overlapping with the cytostomal puncta. Instead, a continuous circle or “necklace of beads” surrounding the parasite was observed as previously reported, sometimes with 1-3 PVM loop extensions in older parasites (4–6, 8, 13, 53, 59, 74). This likely indicates that PTEX does not directly deposit Hb proteases into the cytostome, however this should be more extensively investigated with markers for both the cytostome (Kelch 13) and PTEX (75).

Following uptake into the PV the Hb proteases possibly employ signals within their N-terminal pro-sequences to specify cytostome trafficking. This signal is likely present between aa 84 to 105 of FP2a, given that truncation studies on FP2a show that reporter cargoes with <84 aa of the FP2a N-terminus accumulate at the PPM but 95 aa and 105 aa show partial and complete trafficking to the food vacuole, respectively (35). We also found that the length of the FP2a reporter appears to be important for efficient trafficking to food vacuole as the 190 aa FP2a reporter trafficked more efficiently than the 120 aa reporter which could contribute to more efficient PPM extraction.

In conclusion, we propose a new model for Hb protease trafficking involving PTEX where the early acting FP2a, PM II and possibly other Hb proteases of these families are extracted by the PTEX complex from the PPM and into the PV space. The Hb proteases do not appear to be subsequently translocated across the PVM and beyond into the iRBC since the 120 aa FP2a reporter was not detected in the iRBC or within cargo-associated PVM loops (Fig 8). Furthermore, our FP2a reporter did not degrade in the presence of EQT and proteinase K as the exported GBP130 protein did. We therefore propose that HSP101 could bind to the short 34 aa N-terminal section of FP2a that precedes the TM domain and projects into the PV. The HSP101/FP2a complex could then dock with the rest of PTEX, which may activate HSP101 activity to extract the FP2a reporter into the PV. Alternatively, the intact PTEX complex could directly bind the N-terminal region of FP2a reporter and extract the protease into the PV. The pro-domain of the Hb proteases could then help chaperone the proteases to the lumen of the cytostome via an unknown process, where the proteolytic domains could then face the interior of the Hb containing vesicles that bud off the cytostome (Fig 8). Thereby, PTEX would act as a facilitator of the correct orientation of the Hb proteases for their entry into the cytostome. Although our study has not resolved the complete mechanism of protease transfer into the Hb containing cytostomal vesicles, it could serve as a future starting point to better understand this important process.

## Material and Methods

### Cloning

The *fp2a* gene was appended with a single HA-tag followed by a *glmS* before the 3’ stop codon using CRISPR/Cas9. The construct was designed using a multi-step PCR where sections were amplified from 3D7 genomic DNA and sewn together with the tagging sequence as shown in supplementary data (S2A Fig). The PM II-mScarlet was designed the same way (S7A Fig). Flanks were then ligated into a pBSK bluescript plasmid (Stratagene) and 50 μg of DNA was co-transfected into 3D7 WT (FP2a) or PTEX150-HA*glmS* (PM II-mScarlet) parasites with 50 μg of a Cas9 expressing plasmid (pCas9) under the U6 promoter, containing relevant guide RNA and the human dihydrofolate reductase (hDHFR) cassette (76). The mutated version of the Cas9 plasmid (mpCas9) was prepared for PM II-mScarlet, where the hDHFR was exchanged for a Blasticidin S Deaminase (BSD) drug selection marker since the PTEX150-HA*glmS* line was already WR-resistant. The BSD coding sequence from pEF-Hyp1-Nluc-DH-APEX (59) was amplified in two overlapping fragments so that an internal *BbsI* site could be synonymously mutated to prevent unwanted cleavage prior to insertion of the guide RNA. The first BSD fragment was amplified with BSD_NcoF and BbsI_MutR and the second BSD fragment was amplified with BbsI_MutF and BSD_SacIIR under standard conditions. PCR fragments were then sewn together with BbsI_MutF and BSD_SacIIR and the full-size BSD coding sequence was excised with *NcoI* and *SacII* and ligated into similarly digested pCas9. DNA primers for guide RNAs were chosen according to the list provided by (77) and annealed together prior to ligation into the Cas9 plasmids, which were pre-cut with *BbsI*. The pCas9 was selected for by WR (Jacobs Pharmaceutical) or mpCas9 with Blasticidin S (Life technologies) for 7 days and genomic DNA extracted when parasites came up to confirm correct tagging by PCR. All primers and DNA sequences are listed in S2 Table.

The previously published plasmid pEF-Hyp1-Nluc-DH (53) was used to generate the three different FP2a constructs, pFP2a 120/FP2a 190/ FP2a NT-Nluc-DH-FL. The Hyp1 region of the reporter was removed by excision with *XhoI* and *NcoI* and replaced with the relevant FP2a sequence. FP2a 120 aa and 190 aa contained the first 120/190 aa from the FP2a gene. To generate the FP2a NT, a forward primer was designed after the TM (57 aa of the FP2a sequence) and coupled with the reverse primer created for the 190 aa construct giving a ~120 aa sequence. Plasmids were transfected into the HSP101-HA*glmS* parasite line and expressed episomally under Blasticidin S selection. Primers are listed in S2 Table.

### Parasite culturing

Continuous culture of *P. falciparum* was maintained at 4% haematocrit in human RBCs in AlbumaxII media (RPMI-1640, 25 mM HEPES (Gibco), 367 μM hypoxanthine (Sigma-Aldrich), 31.25 μg/mL Gentamicin (Gibco), 0.5% AlbumaxII (Gibco) and 25 mM NaHCO_3_ (AnalR)) and kept at 37 °C in gas chambers (1% O_2_, 5% CO_2_ and 96% N_2_). Method adapted from Trager and Jensen (78).

### Immunofluorescence assays

For WR trapping experiments, parasite culture was treated with 100 μg/mL heparin (Sigma-Aldrich) to prevent invasion (79) and late schizont stages purified using a 67% percoll (Cytiva, diluted in PBS/RPMI) gradient. Heparin was then washed off and parasites allowed to invade for 4 h. Ring stage parasites were then treated ± 10 nM WR. For BFA assays, parasites were synchronised in 5% sorbitol (Sigma-Aldrich) for 10 min at 37 °C. Late ring/early trophozoite stage parasites were then treated with 18 μM BFA/DMSO or 0.001% DMSO for 5 h and then culture harvested for IFA. For haemozoin crystal experiments, HSP101-HA*glmS* and PTEX150-HA*glmS* parasite lines were sorbitol synchronised at ring stage and treated ± 2.5 mM GlcN at trophozoite stage and harvested the next cell cycle at late ring stage/early trophozoite stage for IFA.

For all assays, parasite culture was diluted in phosphate buffered saline (PBS) and mounted on Poly-L-Lysine (Sigma-Aldrich) treated coverslips (Menzel), fixed in 4% paraformaldehyde/ 0.0075% glutaraldehyde in PBS for 20 min and permeabilised in 0.1% TX-100 (Sigma-Aldrich) with 0.1 M glycine in PBS for 15 min as previously described (80, 81). Samples were blocked for 1 h in 3% bovine serum albumin (Sigma-Aldrich) and subsequently incubated in primary antibodies (overnight) and secondary antibodies (1 h) with 3x 0.02% TX-100/PBS washes in-between. Coverslips were mounted on slides containing mounting media with DAPI (Vectashield) and sealed with nail polish. For the saponin lysis experiment, cells were incubated with ice-cold 0.05% saponin/ PBS on ice for 10 min, pelleted at 50 g for 3 min and washed 3x in 500 μL PBS. Cells were then fixed and probed with antibodies as described above. Images were visualised using the Zeiss Axio Observer Z1 inverted widefield microscope and processed using Image J software. List of antibodies can be found in S3 Table.

Pearson’s correlation coefficients were used to determine co-localisation of Nluc and EXP2. Images were acquired with identical exposure settings and analysis completed on raw images using Fiji software with JACoP plugin. For the saponin lysis experiment, exposure settings were the same for the fluorescent channel of interest for all cells (± GlcN) for direct comparison of signal intensity using raw images. To measure signal only within the parasite, an area was manually drawn around the parasite using the EXP2 signal (PVM) for guidance. To quantify the signal, the integrated density of cells was measured using Image J software and signal corrected by subtracting the background fluorescence signal (cell integrated density – background mean * cell area). Background mean was measured for each field of view used for the analysis, where the average signal of three background areas was subtracted from the cells analysed within the same field of view. For haemozoin crystal experiments, the area was calculated as described for saponin experiments using the DIC channel for guidance and crystals counted as “present” or “absent” completed by two individual counters. All statistical analysis was completed using GraphPad 8 Prism software using Student’s t test with Welch correction. Number of cells analysed and replicates are indicated within each figure.

### Live cell imaging

Heparin synchronised parasite culture was diluted in media 1:1 and evenly distributed over a glass slide and a coverslip placed on top and sealed with wax. Cells were visualised by microscopy as described for IFAs. Analysis was done using Image J where the mScarlet signal was categorised into either ‘dot’ inside the parasite or ‘crescent/circle’ at the parasite periphery by looking at the DIC channel and the Scarlet channel. Three individual counters completed categorisation of cells.

### Growth assays

Sorbitol synchronised trophozoite stage parasite culture was adjusted to 1% haematocrit and 0.3% parasitemia and treated with 0, 0.15 or 1 mM GlcN and subsequently plated in 100 μL aliquots on 96-well plates in technical triplicate. Parasite culture was harvested at trophozoite stage in each cell cycle, for three consecutive cell cycles and stored at −80 °C until all time points had been collected. LDH was then measured as a proxy for parasite growth as previously described (50, 51), where 30 μL of parasite culture was resuspended with 75 μL of malstat reagent (0.083 M Tris, 185 mM lactic acid [adjusted to pH 7.5], 0.17% TX-100, 0.83 mM acetylpyridine adenine dinucleotide, 0.17 mg/mL Nitroblue tetrazolium, 0.08 mg/mL phenazine ethosulphate). Plates were incubated in the dark for 45 min and absorbance measured at 650 nm. Growth was normalised to time point zero (assay set up) and 0 mM GlcN set as 100% growth for each cycle. Statistical analysis was done using Student’s t test with Welch correction in GraphPad Prism 8 for 2 biological replicates.

### Western blotting

Trophozoite stage parasites were lysed in 0.09% saponin (Kodak) in PBS containing protease inhibitors (PI, Roche) on ice for 10 min. For knockdown experiments and FP2a reporter expression experiments, heparin synchronised (4 h invasion window) trophozoite stage parasites were treated for one cell cycle (trophozoite to trophozoite) with GlcN (Sigma-Aldrich) and parasite pellet extensively washed in PBS containing protease inhibitor cocktail (PI, Roche) after saponin lysis. For other experiments, parasites were sorbitol synchronised. Parasite pellets were diluted in 1x sample buffer (6x: 0.3 M Tris (Astral Scientific, 60% v/v glycerol (Astral Scientific), 12 mM EDTA (Sigma-Aldrich), 12% SDS (Sigma-Aldrich), 0.05% bromophenol blue (BioRad)), sonicated 3x cycles (Bioruptor pico, 30 sec on/30 sec off) and reduced in 100 mM Dithiothreitol (DTT, Sigma-Aldrich) prior to fractionation on 3-12% Bis-Tris gels (Invitrogen). Proteins were transferred to nitrocellulose membranes (iBlot, 20V, 7 min), blocked in 1% casein/PBS (Sigma-Aldrich) for 1 h and incubated in primary antibodies (S3 Table) overnight and in secondary antibodies for 1 h (Invitrogen) in the dark. For chemiluminescence detection of chicken anti-FLAG and rat anti-RFP (to detect mScarlet), the membrane was additionally incubated for 5 min in SuperSignal substrate (Pierce). Blots were visualised and densitometry analysis performed using the Odyssey LI-COR system. Graphs and statistical analysis, Student’s t test with Welch correction or simple linear regression, were completed using GraphPad 8 Prism Software. Full-length western blots in this study can be found in supplementary information (S8 – S18 Figs).

### Co-immunoprecipitation assays

Sorbitol synchronised ring stage parasites were treated ± 10 nM WR for ~16 h and subsequently harvested at trophozoite stage (~28-32-hpi) via magnet purification (LS, Militeyni). The iRBC pellet was resuspended in 20x pellet volume of 1x modified radioimmunoprecipitation (RIPA) buffer (1% TX-100, 0.1% SDS, 10 mM Tris [adjusted to pH 7.5], 150 mM NaCl) (56) containing PI. Cells were lysed via 2x cycles of freeze/thawing, pelleted at 16,000 g for 10 min at 4 °C and resultant supernatant was transferred to a new tube. Two different methods were used, either HA or IgG IP assays:

1. For HA IP assays, 250 μL of supernatant (~2.5 mg of protein) was diluted in 750 μL of lysis buffer and 50 μL of diluted supernatant transferred to a new tube and used as assay input. Diluted supernatant was incubated overnight with 50 μL of pre-washed 50/50 anti-HA agarose bead slurry (Sigma-Aldrich, diluted in lysis buffer)
2. For IgG IP assays, 200 μL of supernatant (2 mg of protein) was diluted in lysis buffer to make 1 mL total volume and 50 μL of diluted supernatant transferred to a new tube and used as assay input. The diluted supernatant was incubated with IgG overnight. The next day, IgG assay samples were incubated with 100 μL of pre-washed 50/50 Protein Sepharose A slurry (Sigma-Aldrich, diluted in lysis buffer) for 1 h. Samples for both HA and IgG assays were then passed through a Micro-Bio spin column (Bio-Rad) and eluted in 50 μL 1x sample buffer for 5 min at room temperature. Input was resuspended in 10 μL of 6X sample buffer and both input and elution were reduced (HA IP) by addition of in 100 mM DTT or kept non-reduced (IgG IPs) prior to SDS-PAGE and western blotting.

### Proteinase K protection assay

Sorbitol synchronized trophozoite stage parasites (~24-hpi) were enriched via magnet purification and parasite pellet used immediately for proteinase K protection assays as previously described (82) but with a few modifications. 20 μL of the iRBC pellet was resuspended in 80 μL of PBS + PI supplemented with 1.6 μg of recombinant EQT (made in-house) and incubated for 10 min at 37 °C. Samples were centrifuged at 1,000 g for 3 min at 4 °C and supernatant transferred to a new tube and kept on ice. The pellet was subsequently washed 3x in PBS and divided equally between four tubes. Samples were pelleted and supernatant discarded. 100 μL PBS was added to tube 1, 100 μL PBS + proteinase K (20 μg) to tube 2, 100 μL 0.03% saponin in PBS + proteinase K to tube 3 and 100 μL of 0.25% v/v TX-100 in PBS + proteinase K to tube 4. All tubes were incubated on ice for 30 min. To stop the proteinase K activity and precipitate proteins, 11.1 μL of trichloroacetic acid (TCA, AnalR) was added to each tube, resuspended thoroughly and incubated on ice for 10 min. Samples were centrifuged at 16,000 g for 20 min at 4 °C. Supernatant was discarded and 500 μL of 100% ice-cold acetone (Merck) was added to the pellet and centrifuged at 16,000 g for 10 min at 4 °C. Supernatant was discarded and pellet left to air-dry. The dried pellet was resuspended in 100 μL of 1x sample buffer, and 26 μL of 4x sample buffer was added to EQT supernatant. All samples were reduced in 100 mM DTT and prepared for western blotting.

### Metabolite extraction and analysis

Sorbitol synchronised parasite culture with ~10% trophozoite stage parasites (~24 hpi) were treated with 0, 0.15 or 1 mM GlcN plus heparin (100 μg/mL) to prevent parasite invasion. At late schizont stage the heparin was removed and parasite culture concentrated to 8% haematocrit and left shaking (85 RPM) at 37 °C for 4 h to allow parasites to invade new RBCs. Culture was then sorbitol synchronised and parasite pellet placed back into culture at 1% haematocrit at 37 °C. GlcN concentration was maintained throughout all steps, except during sorbitol treatment. PTEX150-HA*glmS* parasite culture was harvested at 18, 24 and 30-hpi or at a single time point 24-hpi including additional 3D7 WT control. For metabolite extraction, samples were prepared in three technical replicates, each replicate containing 1 mL warm media and 1×10^8 cells. All 12 tubes (0, 0.15, 1 mM GlcN and uRBCs) were incubated at 37 °C. Technical triplicates were processed together for the following steps at 4 °C. Samples were centrifuged at 12,000 g for 30 s, supernatant removed, and pellet resuspended in 1 mL ice-cold PBS. Samples were centrifuged as before, and final pellet was resuspended in 200 μL of 80% acetonitrile (Merck, in Milli-Q water). The samples were centrifuged at 18,000 g for 5 min, supernatant transferred to fresh tubes and stored at −80 °C until processed by mass spectrometry as previously described (83).

Polar metabolite detection was performed on an Agilent 6550 Q-TOF mass spectrometer operating in negative mode. Metabolites were separated on a SeQuant ZIC-pHILIC column (5 μM, 150 X 4.6 mm, Millipore) using a binary gradient with a 1,200 series HPLC system across a 45 min method using 20 mM ammonium carbonate (pH 9) and acetonitrile, as described previously (83). The scan range was 25-1,200 m/z between 5 and 30 min at 0.9 spectra/s. An internal reference ion solution was continually run (isocratic pump at 0.2 mL/min) throughout the chromatographic separation to maintain mass accuracy. Other liquid chromatography parameters were: autosampler temperature 4 °C, injection volume 5-10 μL and data was collected in centroid mode with Mass Hunter Workstation software (Agilent). Raw Agilent.d files were converted to mzXML with MSconvert and analysed using the Maven software package (84). Following alignment, metabolite identification was performed either with exact mass (<10 ppm) or retention time matching to authentic standards (approximately 150 in-house metabolite standards).

## Acknowledgements

We thank the Australian Red Cross for the human RBCs used in this study and Jacobus Pharmaceutical for providing the WR. We thank Cat Nie for technical assistance with troubleshooting of the initial metabolomics experiment, analysis and sample preparation. We thank Paul R. Sanders for technical assistance during the preliminary work of this project. We also thank Leann Tilley and Matthew Dixon for gifting the CRT, ERC and GAPDH antibodies and the Malaria Experimental Genetics 2018 course for providing the pCas9 plasmid.

## Author contributions

T.K.J, roles: Conceptualization, Data Curation, Formal Analysis, Investigation, Methodology, Validation, Visualization, Writing – Original Draft Preparation, Writing – Review & Editing. B.E., roles: Conceptualization, Investigation, Methodology. S.C., roles: Formal analysis, Investigation, Methodology. M.G., roles: Formal Analysis, Investigation, Methodology. S.C.C., role: Investigation. M.G.D., role: Formal Analysis. M.P.S., role: Formal Analysis. M.M., roles: Resources, Funding acquisition. H.E.B., roles: Conceptualization, Data curation, Funding acquisition, Methodology, Project Administration, Resources, Supervision, Visualization, Writing – Review & Editing. B.S.C., roles: Conceptualization, Data curation, Funding acquisition, Methodology, Project Administration, Resources, Supervision, Writing – Review & Editing. P.R.G., roles: Conceptualization, Data curation, Funding acquisition, Investigation, Methodology, Project Administration, Resources, Supervision, Writing – Review & Editing

## Supporting information

**S1 Fig. Metabolite heat maps.**

(A) Heat map corresponding to S1 Table for PTEX150-HA*glmS* metabolomics time course (18, 24, 30-hpi) experiment. (B) Heat map corresponding to S1 Table for 3D7 and PTEX150-HA*glmS* metabolomics experiment. Numbers represent metabolites listed in S1 Table, where Hb peptides have been highlighted in bold. Fold change for both heat maps was calculated using the following formula =log2(GlcN treated/untreated).

**S2 Fig. Generation and characterisation of the FP2a-HA*glmS* parasite line.**

(A) CRISPR-Cas9 was used to append the *fp2a* gene with a HA tag and a *glmS* riboswitch. (B) PCR was used to confirm the correct integration of the tag to the target gene, where 3D7 WT was used as a negative control for integration. The position of PCR primers used to confirm integration are shown in panel A, where FInt, is the forward PCR integration primer. F, forward primer. glmS_R, reverse primer. (C) Correct size and level of FP2a knockdown was confirmed using western blotting. >90% of FP2a was knocked down after one cell cycle of GlcN treatment. P stands for pro FP2a and M stands for mature FP2a. The HA-tag adds approximately 3 kDa to the target protein. The mark on the right-hand side of the blot is an artefact. Mouse anti-HA detects FP2a-HA*glmS* and rabbit anti-HSP70-1 was used as a loading control. Blot is representative of 3 biological replicates. (D) Immunofluorescence assays where mouse anti-HA detects target protein and rabbit anti-EXP2 was used as a PVM marker. The images indicate that FP2a displays diffused localisation in the parasite cytoplasm, sometimes with a concentrated puncta at the parasite periphery, likely representing the cytostome (white arrows). The HA antibody signal against FP2a-HA*glmS* partly overlaps with rabbit anti-CRT which labels the food vacuole membranes. Food vacuole lumen, indicated by the dark haemozoin crystals in the DIC images, are not well labelled for FP2a-HA*glmS* likely due to the HA epitope tag being proteolytically degraded in the food vacuole. Scale bars= 5 μm. (E) Multi-cycle LDH growth assays were performed over 3 consecutive cell cycles. No significant growth defect was observed for FP2a or 3D7 WT when treated with different concentrations of GlcN. T = number of cycles. Statistics were completed using Student’s t test with Welch correction. Graphs show 2 biological replicates completed in technical triplicate. Error bars= SD. Data for each parasite line was normalised to 0 GlcN treatment for each time point.

**S3 Fig. Knockdown of PTEX150 and HSP101 results in build-up of full-length Hb inside the parasite and reduced haemozoin crystal formation.**

(A) Simple linear regression analysis was performed on protein expression and Hb build-up from western blots presented in Fig 2B for PTEX150-HA*glmS*, HSP101-HA*glmS* and FP2a-HA*glmS*. All parasite lines showed significant regression slope, where P values are shown in each graph along with R^2^. The 6 blue dots on the graphs represent 3 biological replicates for 0.15 and 1 mM GlcN for protein expression (x-axis) plotted against the mean for 3 biological replicates for Hb build-up (y-axis). (B) Both PTEX150-HA*glmS* and HSP101-HA*glmS* trophozoite stage parasites were treated ± 2.5 mM GlcN for one cell cycle and harvested for IFA. Haemozoin crystals in the DIC channel were counted (present or absent). Images are representative of 3 (PTEX150) or 2 (HSP101) biological replicates. (C) Both PTEX150 and HSP101 knockdown experiments shown in panel B resulted in significantly less crystal formation compared to untreated cells when using Student’s t test with Welch correction. (*) Indicates P= 0.0247 (PTEX150) and P=0.0277 (HSP101). Error bars = SD from 2 individuals counting. (D) Area of the parasites analysed in panel C (completed as described in Fig 2E) showed significant difference in size for GlcN treated PTEX150 parasites compared with untreated indicating parasite growth was affected but no significant difference was observed for HSP101 knockdown. Middle line represents mean and error bars=SD. (****) Indicates P=<0.0001. Each dot on the graph represents one cell analysed.

**S4 Fig. Western blot time course indicates that the 120 and 190 aa FP2a reporters are expressed similarly from 24-40-hpi while the NT reporter is degraded as the parasite matures.**

Synchronous (4 h invasion window) parasite cultures for the three FP2a reporter lines were divided in four and harvested via saponin lysis at four sequential time points: 24-28 hpi, 28-32 hpi, 32-36 hpi and 36-40 hpi. Chicken anti-FLAG and rabbit anti-Nluc were used to visualise the FP2a reporters, mouse anti-HA to visualise the HSP101-HA*glmS* (parental line) and rabbit anti-HSP70-1 was used as a loading control. The expected size of each reporter is indicated with (*), where the cleavage of Nluc from the 120 and 190 aa reporters was observed in each time point (lanes 2, 3, 5, 6, 8, 9, 11 and 12), likely due to cleavage upon entry into the food vacuole. This cleavage of Nluc was not observed for the NT reporter (lanes 1, 4, 7 and 10), which does not enter the food vacuole. The NT reporter was also degraded more, and the expression of the full-length reporter diminishes as the parasite matures (lanes 1 and 4 vs. lanes 7 and 10), whilst expression of the 120 and 190 aa reporters remains stable across each time point. These data indicate that the optimal time point to study these three reporters was in the range of 24-32 hpi, as indicated in bold. This blot represents 2 biological replicates.

**S5 Fig. The 120 aa and the 190 aa FP2a reporters interact with PTEX150 and EXP2.**

Immunoprecipitation assays with parasites treated with ± 10 nM WR were completed on the four reporter lines using Protein Sepharose A where (A) PTEX150 or (B) EXP2 protein specific antibodies were incubated with parasite lysate to Co-IP interacting proteins. Both 120 aa (line 13) and 190 aa (line 15) showed stronger association with (A) PTEX150 and (B) EXP2 in –WR treatment as determined by densitometry (S6 Fig). The Hyp1 reporter showed reverse association to FP2a reporters and NT showed some background signal in both conditions. (A) PTEX150 and (B) EXP2 both co-precipitated with other PTEX components (lanes 9 – 16). Chicken anti-FLAG was used to probe for the reporter, mouse anti-HA for HSP101 and rabbit 741 for PTEX150. Both (A) and (B) blots represent 3 biological replicates. The asterisks indicate stronger signal when comparing the ± WR treatment for respective reporter measured by densitometry of 3 biological replicates.

**S6 Fig. Densitometry measurements for PTEX150 and EXP2 IPs.**

(A) The interaction of reporters with PTEX150 (r942) and EXP2 (r1167) presented in S5 Fig A was graphed, where FLAG elution was adjusted to input. Each dot represents 1 biological replicate. (B) The interaction of reporters with PTEX150 (r942) and EXP2 (r1167), where the fold difference of the FLAG elution/input was adjusted to untreated (-WR). Error bars= SD from 3 biological replicates.

**S7 Fig. Establishment of PM II mScarlet in a PTEX150-HA*glmS* background.**

(A) The *pm ii* gene was C-terminally tagged with mScarlet as described for FP2a in S2 Fig A and introduced into a PTEX150-HA*glmS* background. (B) Correct integration of mScarlet to PM II was confirmed via PCR, where genotyping primers are displayed in panel A.

**S8 – S18 Figs. Full-length western blots presented in this study.**

